# Topological conditions for propagation of spatially-distributed neural activity

**DOI:** 10.1101/2021.11.30.470616

**Authors:** Àlex Tudoras, Alex D. Reyes

## Abstract

An important task of the nervous system is to transmit information faithfully and reliably across brain regions, a process that involves the coordinated activity of a relatively large population of neurons. In topographically organized networks, where the entering and exiting axons of neurons terminate in confined areas, successful propagation depends on the spatial patterns of activity: the firing neurons in a presynaptic or source layer must be located sufficiently close to each other to ensure that cells in the postsynaptic or target layer receive the requisite number of convergent inputs to fire. Here, we use principles of topology to define the conditions for transmitting information across layers. We show that simplicial complexes formed by source neurons can be used to: 1) determine whether target neurons receive suprathreshold inputs; 2) identify neurons within the active population that contribute to firing; and 3) discriminate between single and multiple active clusters of neurons.

## Introduction

Many sensory systems use topography as a substrate for preserving information across brain regions. Auditory, visual, and somatosensory representations rely on maps -tonotopy, retinotopy, somatotopy- that are formed in peripheral organs and maintained faithfully through brainstem, thalamus, and cortex [1, 2, 3]. The propagation of signals across the successive processing stages generally involves the combined and coordinated activities of a relatively large population of neurons. Though the conditions leading to successful propagation has been examined extensively [4, 5, 6, 7, 8, 9, 10, 11, 12], the focus has been primarily on the temporal features and less so on the spatial aspects of neuronal firing.

In topographically-organized systems, the axon from a neuron in one network (henceforth termed source layer) branches out to contact several neurons in a downstream network (target layer) [3, 13]. The axon collaterals are not scattered uniformly throughout the target network but tend to contact neurons within a confined area (henceforth termed synaptic field) [3, 13, 14, 15] (Fig. 1a). Consequently, when a cluster of neurons in the source layer become active, their associated synaptic fields overlap (Fig. 1b*i*). Target neurons will fire if they receive a minimum number of convergent excitatory inputs that raise their membrane potential past the threshold for action potentials.

**Figure 1:**
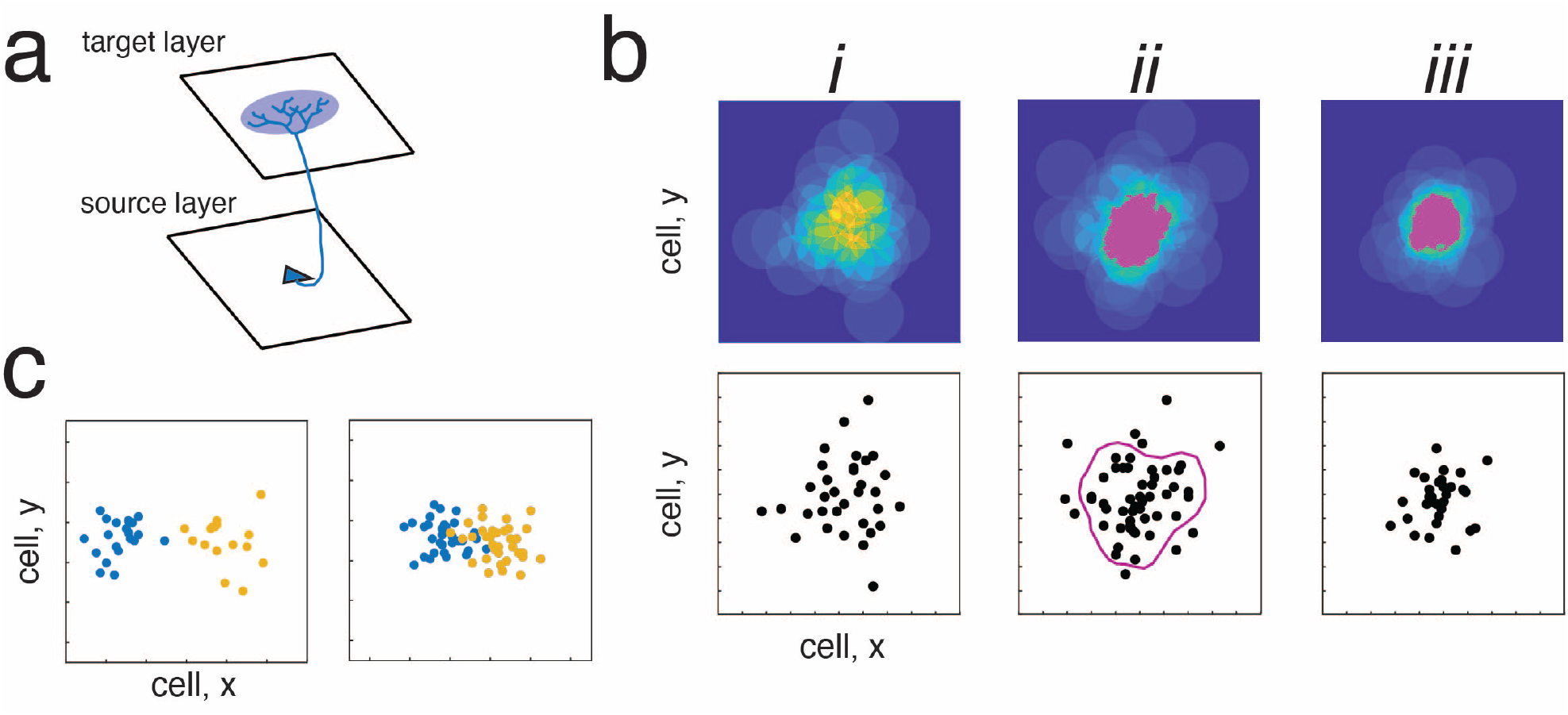
Relation between active neurons and synaptic fields. *a*, synaptic field generated in the target layer by a neuron in the source layer. ***bi***, composite synaptic field (top) generated by a population of active neurons in the source layer (bottom). The overlap in synaptic fields in insufficient to produce suprathreshold inputs. ***ii***, suprathreshold inputs are generated by increasing the number of active cells in the source layer (magenta area, top). The extensive overlap in synaptic fields were generated by source neurons enclosed by the magenta circle (bottom). ***iii***, suprathreshold inputs generated when the spatial distribution of active neurons was narrowed. ***c***, two populations of neurons (blue, orange) spaced widely apart (left) and sufficiently close to each other for the two populations to intermix (right)

Because of the limited extent of synaptic fields, the magnitude of the input to target neurons depends on the spatial distribution of active neurons in the source layer. If the active neurons are spaced too widely apart (Fig. 1b*i*), the overlap of synaptic fields in the target layer is insufficient to evoke firing. To increase overlap, the number of active neurons within an area could be increased (*ii*) or, alternatively, the spatial distribution of a comparable number of active neurons could be reduced (*iii*). The effect in both cases is to increase the local density of active neurons.

From a coding perspective, it will be important to develop criteria for identifying and categorizing the spatial patterns that are conducive for signal propagation across layers. At the microscopic level, inhomogeneities in the distribution of active neurons may mean that not all neurons contribute equally to firing in the target layer. In Fig.1B*ii*, for example, only neurons enclosed by magenta circle (bottom) are able to generate sufficient overlapping synaptic fields in the target layer that exceeds threshold for firing (magenta region in top); other active neurons away from the center do not contribute and their associated synaptic fields remain subthreshold (light blue regions). At a more macroscopic level, simultaneous stimuli or a stimulus with multiple components can lead to complex spatial patterns of activity. For example, the two clusters of neurons in Fig. 1C, (orange, blue) are easily distinguishable from each other when they are far apart. However, if the clusters are too close, they begin to intermix and it becomes difficult to determine reliably whether one or two activity patterns are propagated to the next layer.

Here, we use topological analyses to describe quantitatively the spatial patterns in the source layer that generate suprathreshold inputs in the target layer. We show that the conditions for successful propagation can be related to the dimensions of simplicial complexes formed from the active source neurons. We then use Betti numbers to examine global and local spatial features and to distinguish between single and multiple clusters of active neurons.

## Results

In the following sections, we first define formally the source and target neural spaces that will be used for the analyses. We then quantify the spatial firing patterns of source neurons that are conducive for generating suprathreshold events in the target layer by constructing Čech complexes and identifying ‘functional’ simplices.

### Neural Space

Biological neural networks are composed of neurons that are embedded in and separated by extracellular matrix. Because the neurons are the functional units, the neural space and associated sensory representation are essentially discretized [16]. To facilitate formal analyses, we disregard the extracellular matrix and construct the neural space with a grid-like architecture. The network consists of a pair of two dimensional sheets of neurons in a feedforward configuration where neurons in a ‘source’ layer (*S*^*α*^) send axons to neurons in a ‘target’ layer (*S*^*β*^) (Fig. 1a). The partitioning of neural space into units of neurons described previously [16] is extended to 2 dimensions (see Appendix). Visually, the neural space is a ‘half-open’ box with the bottom and left edges closed and the top and right edges open (Fig. 2A). Individual neurons (inset) are also ‘half-open squares’ with width Δ*h*. Defining the space in this manner is convenient since there are no ‘gaps’ between neurons, reflecting the absence of the extracellular matrix.

**Figure 2:**
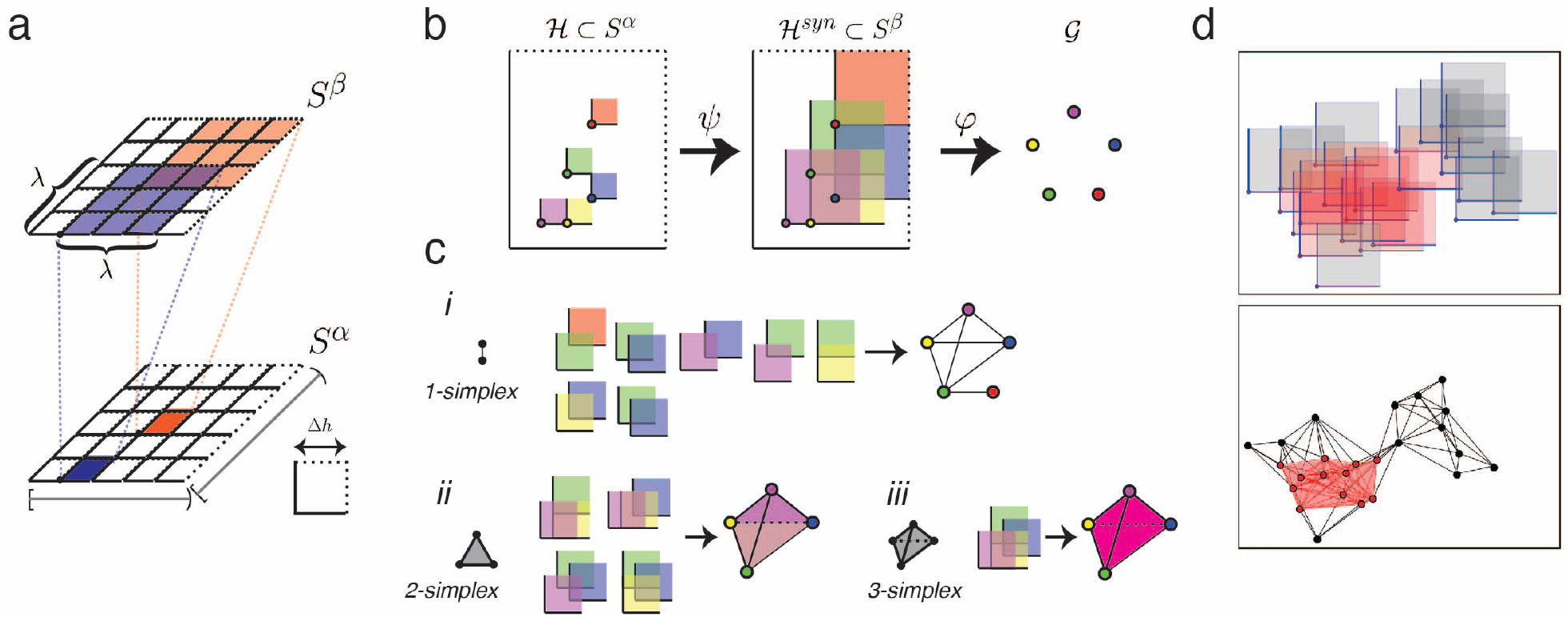
Construction of simplices. *a*, hypothetical two layer network of neurons. Layers are half-open rectangles that are partitioned into neurons, each of which is a half-open square box with length Δ*h* (bottom, inset). Neurons in *layer S*^*α*^ generate rectangular synaptic intervals in *layer S*^*β*^ with dimension *λ*_*x*_ × *λ*_*y*_. ***b***, Relation between elements of source layer (*s*^*α*^), target layer (*s*^*β*^), and graph (𝒢). Neurons in the source layer *S*^*α*^ (half-open boxes) generate overlapping synaptic fields in the target layer *S*^*β*^ via the mapping *ψ*. The corner points are mapped to a graph 𝒢 via the mapping *ϕ*. ***c, i***, 1-simplices are formed when 2 synaptic fields overlap. The graph consists of connected points that form unfilled triangles (right). ***ii***, 2-simplices are formed 3 synaptic fields overlap. The triangles are filled but the tetrahedron is hollow (right). ***iii***, 3-simplex formed 4 overlapping synaptic fields. The resulting tetrahedron is solid. ***d, bottom***, Cloud of active neurons and the functional *k*−simplices *k* ≥ *n*_*θ*_ − 1, *n*_*θ*_ = 8 shown with red vertices and lines and shaded; the non-functional *k*−simplices *k < n*_*θ*_ − 1 shown in black. ***top***, synaptic fields generated in target layer.

The individual neurons are uniquely identified by the point at their lower left corner (termed ‘corner point’). Henceforth, neurons will be designated as *h*_*xy*_ = [*x, x* + Δ*h*) × [*y, y* + Δ*h*). Letting the sets containing the starting points of intervals in both axes be χ = {*x*_*i*_ = *i* · Δ*h* | 0 ≤ *i < N*_*x*_} and 𝒴 = {*y*_*j*_ = *j* · Δ*h* | 0 ≤ *j < N*_*y*_}, the set of cells can then be expressed as:

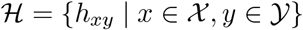

A topology *τ*_*S*_ for neural space *S* can be defined with ℋ as a basis (see Appendix).

The synaptic field generated in the target layer *S*^*β*^ is also a half-open square that encompasses *n*_*λ*_ × *n*_*λ*_ neurons (Fig. 2A, top). Each synaptic field can be identified by the location of the neuron in the lower left corner and expressed as:

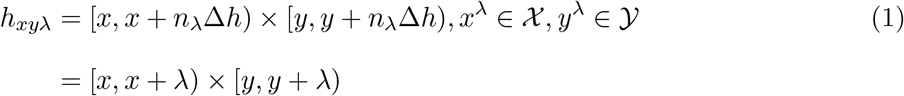

where χ^*λ*^ = {*x*_*i*_ | 0 ≤ *i < N*_*x*_ − *n*_*λ*_} ⊂ χ ; 𝒴^*λ*^ = {*y*_*j*_ | 0 ≤ *j < N*_*y*_ − *n*_*λ*_} ⊂ 𝒴 to ensure that the synaptic fields are within neural space.

A mapping *ψ* from individual neurons the source space (*S*^*α*^) to synaptic fields in the target space (*S*^*β*^) can be described as follows. For simplicity, we take *S*^*β*^ to be a copy of *S*^*α*^ so that neurons and synaptic intervals can be identified by the same coordinates (*x, y*) ∈ χ × 𝒴 (see Appendix for a more general form). The mapping, with (*x, y*) restricted to χ^*λ*^ × 𝒴^*λ*^, is then:

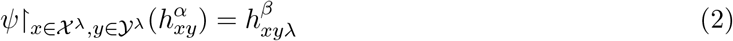

### Simplicial complexes underlying synaptic fields

As stated in the Introduction, the successful propagation of signals across networks depends on the spatial arrangements of active neurons in the source layer. In the following, we will use simplices to describe the spatial distributions of source neurons that lead to firing of target neurons.

The mapping *ψ* defined above will be restricted to cells in *S*^*α*^ that become active during a stimulus. Letting the coordinates of these active cells be the set 𝒜 ⊂ χ^*λ*^ × 𝒴^*λ*^, the map from active neurons in the source layer to synaptic fields in the target layer is:

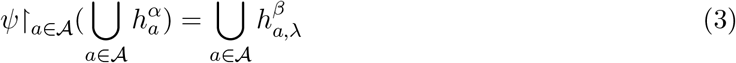

A lemma in the appendix shows that *ψ* is a homomorphism.

Because of topographical organization, the activated source neurons are clustered within a confined area. The resulting synaptic fields in the target layer will overlap (or more formally, intersect), the extent of which is determined by *λ*. The intersecting synaptic fields can be used to construct the simplicial Čech complex in *S*^*β*^, and by extension in *S*^*α*^. Intuitively, as will be shown below, the Čech complex captures the relationship between synaptic overlap and the spatial distribution of source neurons and extent of the synaptic field.

Letting A be the set containing the corner points of synaptic fields generated by active neurons, then the Čech complex is:

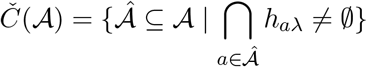

Theorem 1 below states that for the discretized neural space, the Čech complex constructed with rectangular synaptic fields can be made identical to that constructed with the more standard 2 dimensional balls. The rectangular synaptic fields permit precise calculation of the number of neurons in *S*^*α*^ that receive suprathreshold inputs and allows for rapid calculation of the Čech complexes (see Methods). Importantly, the intersection of the synaptic fields is consistent with the topology of the neural space and encompasses an integral number of neurons. In contrast, the intersections of circular synaptic fields in a continuous neural space could fall in regions with no neurons.

#### Theorem 1.

Let *S* ⊂ ℝ^2^ be the neural space that is partitioned into individual square neurons each of width *h* and square synaptic fields with corner points (*a, b*) = (*n*_*a*_ · *h, n*_*b*_ · *h*) ∈ *S* ⊂ ℝ^2^ of length *λ* = *n*_*λ*_ · *h* with *n*_*a*_, *n*_*b*_, *n*_*λ*_ ∈ ℤ, *n*_*λ*_ 0 and *h* ∈ ℝ^+^. If 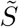 is a space consisting of balls with centers located at 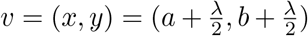 with fixed radii 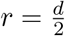 such that 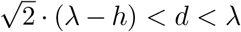, then ∀*I* = {*i*_1_, …, *i*_*n*_}, synaptic fields 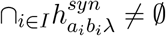 in 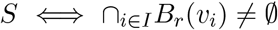.

### Corollary

Under the conditions described in Theorem 1, the Čech complex generated in 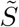 is identical to the one generated from the intersections in the discretized neural space, with the same number of *k*-simplices *S*.

The Čech complex contains simplices of various dimensions depending on the number of nonempty intersection of synaptic fields. The corner points of the synaptic fields can be mapped to vertices in the graph G to obtain the abstract simplices (see Appendix). For example, an edge (1-simplex) is formed when the (Chebyshev) distances between 2 corner points of the synaptic fields is lower than *λ*, signifying overlap (Fig. 2B*i*). Similarly, a filled triangle (2-simplex) and solid tetrahedron (3-simplex) requires non-empty intersection between, respectively 3 and 4 synaptic fields (Fig. 2B*ii, iii*). In general, a (*n* − 1)-simplex has vertices that correspond to the set of corner points of *n* synaptic fields with non-empty intersection.

### Functional simplices

Although there can be many neurons in the source layer that become active, only a subset may actually contribute to firing of neurons in the target layer. A target cell will fire if it receives sufficient convergent inputs (*n*_*θ*_) from the source layer, or equivalently, if the cell is at location where *n*_*θ*_ synaptic fields intersect. This corresponds to the (*n*_*θ*_ − 1)-simplices in *Č*(𝒜). Hence, a large *n*_*θ*_ means that many source neurons need to be sufficiently close to each other to generate significant synaptic overlap; the geometric relationship between these neurons is captured by higher dimensional simplices. We prove in the appendix that:

#### Theorem 2.

In topographically-organized systems, a neuron in the target layer *S*^*β*^ will receive a sufficient number (*n* ≥ *n*_*θ*_) of inputs to fire if there is a (*n* − 1)-simplex in the source layer *S*^*α*^.

The connected points in Fig. 2D (bottom) show the simplicies formed using rectangular synaptic fields (top). The red-filled neurons and shaded region encompasses these functional *k*−simplices with *k* ≥ *n*_*θ*_ −1 = 7 in the source layer. The black-filled neurons are the vertices for lower dimension simplices, indicating that they are spaced too widely to produce the requisite number of overlapping synaptic fields. These active neurons would not lead to firing in the target layer and are considered ‘non-functional’ in this scenario. *k*-simplices with *k > n*_*θ*_ − 1 means that inputs to some target cells exceed threshold level. Though the analyses treats neurons as binary units, the amount that *k* exceeds *n*_*θ*_ − 1 could be interpreted as an increase in the firing rate of target neurons.

Formally, the functional Čech complex 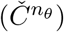 is a subset of the complete Čech complex with the following filtration by dimension of the simplices:

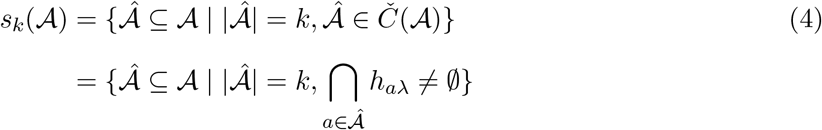

Thus, the components of *Č*(𝒜) that are part of a *k*−simplex, *k* = (*n*_*θ*_ −1) compose the following filtered Čech complex

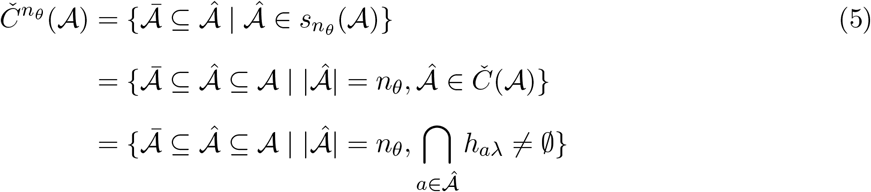

The presence of *k* = *n*_*θ*_ − 1 Čech complex in the source layer means that at least one neuron in the target layer receives threshold inputs and conversely, a target neuron receiving a threshold input means that there is at least one *n*_*θ*_ − 1 simplex in the source layer.

Equation 5 indicates that further quantification of the relation between the Čech complex and synaptic fields will be difficult. First, the number of neurons (vertices) that participate in functional simplices could be the same under different conditions. Second, increasing *n*_*θ*_ raises the minimum number of synaptic fields that must have non-empty set intersection, effectively decreasing the number of target neurons receiving threshold inputs. Fig. 3 shows the Čech complex and associated synaptic fields as *n*_*θ*_ is increased from 2 to 6. The complexes from *n*_*θ*_ = 2 to 4 are identical, but the areas (yellow) receiving threshold inputs shrinks. At *n*_*θ*_ = 5, 6, a few edges and vertices are eliminated and the activated areas decrease further.

**Figure 3:**
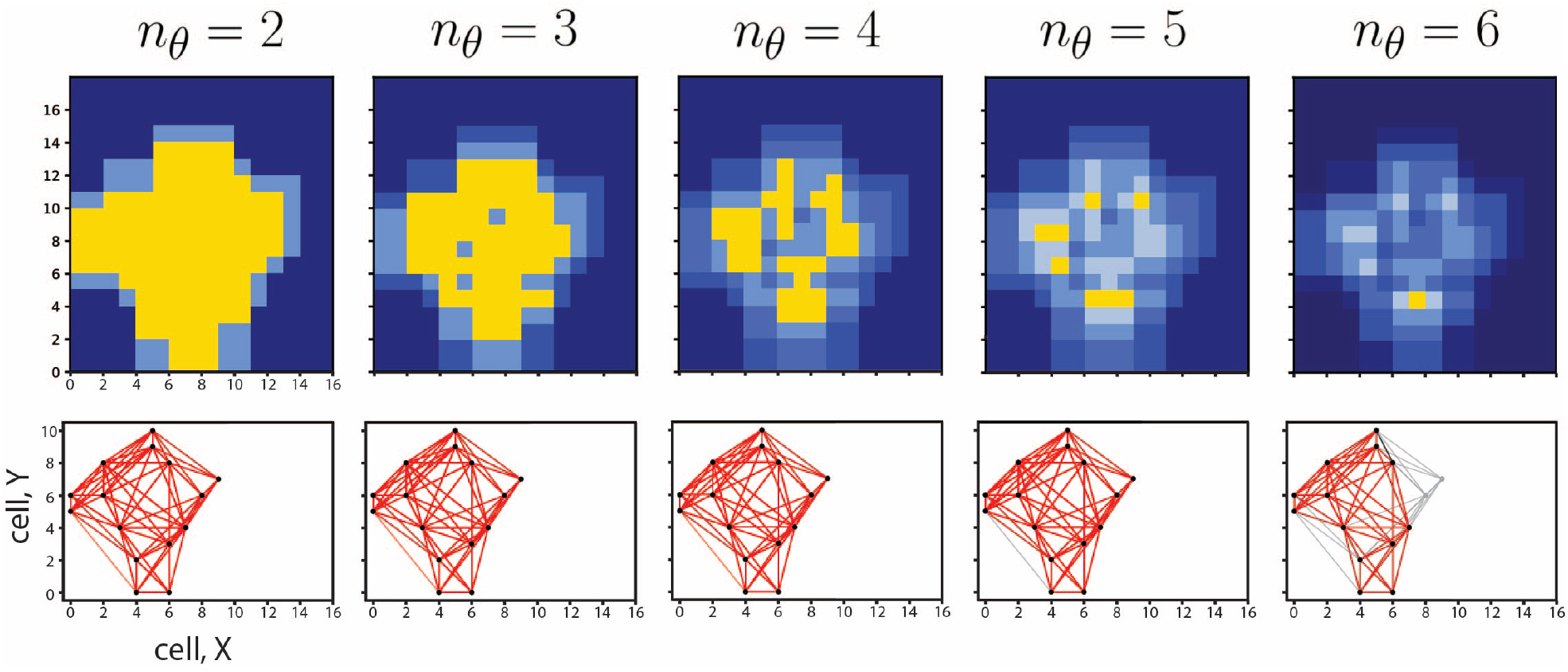
Filtration of Čech complexes by n_θ_, bottom, Čech complex of a cloud of active neurons obtained with (from left to right) *n*_*θ*_ = 2, 3, 4, 5, 6. Gray lines indicate components that do not belong to functional simplices. ***top***, corresponding synaptic fields. Yellow regions indicate suprathreshold input to cells.

### Relation between simplices and synaptic fields

The simplices provide information about the required spacing of active neurons in the source layer for activating cells in the target layer. Given an estimate of *n*_*θ*_ and the spread of the synaptic field *λ*, it is theoretically possible to predict from the dimension of simplices whether a population of active cells will produce sufficient overlap in synaptic field to fire cells in the target layer.

To examine the relation between spacing of active neurons, the resulting simplices, and synaptic fields, we simulated a network with a fixed number of neurons and varied the spatial distribution (Fig. 4). The location of a fixed number of active cells were drawn from a 2 dimensional Gaussian distribution and the spread was adjusted by changing the standard deviation (Fig. 4A, see Methods). As the distribution narrowed (decreasing *σ*), the number of *k*-simplices increased (Fig. 4B, left). The lower dimension 1- and 2-simplices were present even for broad distribution (*σ <* 10); the higher dimension simplices appeared in succession with decreasing *σ* (inset). The number of cells receiving threshold inputs (*n*_*θ*_ = 5) increases with decreasing *σ* (Fig. 4B, right) and follows the initial rise of the 4-simplex (black in B, left). Similar observations were made when *σ* was kept constant and the number of active source cells was increased (Fig. 4C).

**Figure 4:**
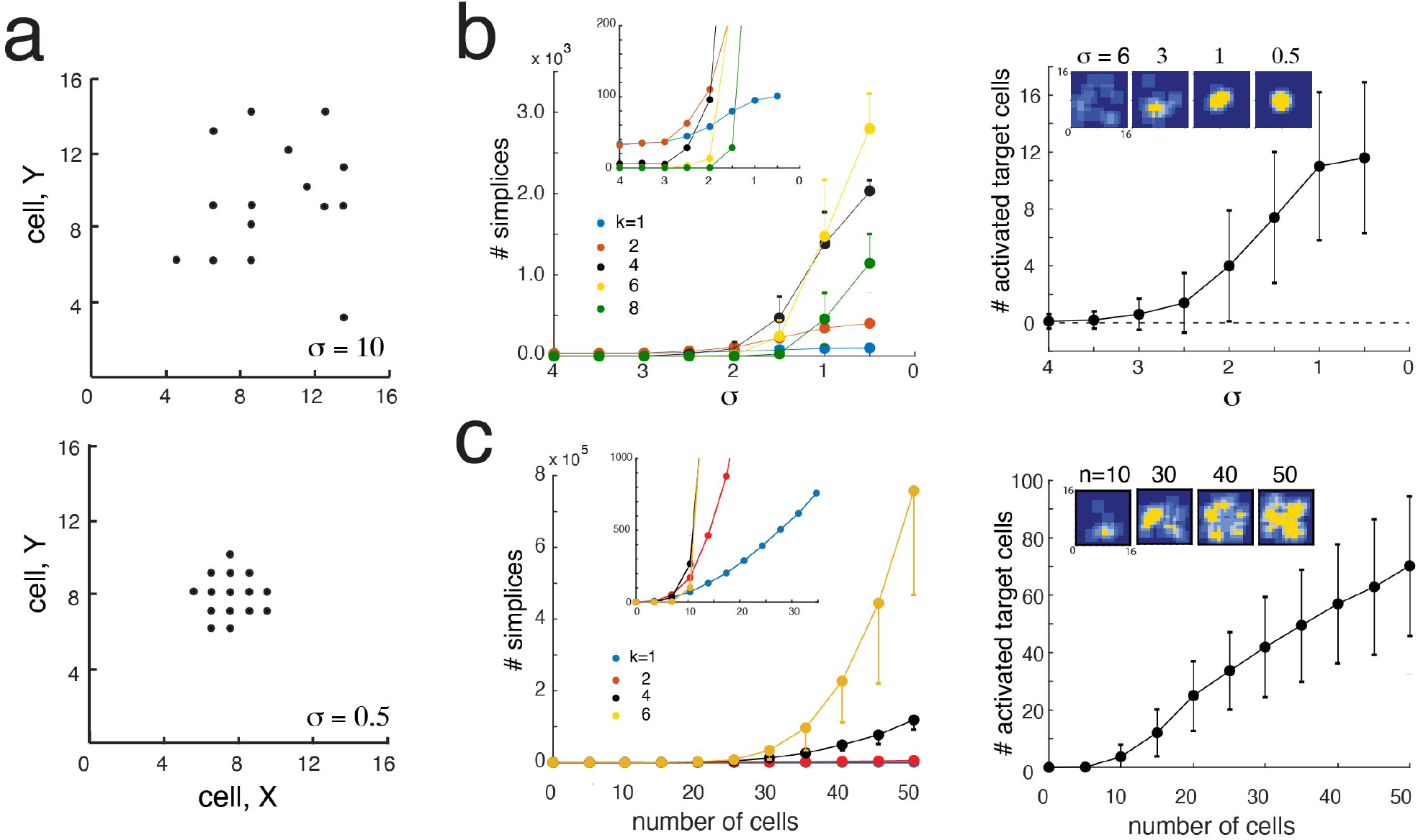
Dependence of simplices on firing cell distribution. ***a***, examples of 15 active source cells broadly (left) and narrowly (right) distributed. Location of cells were drawn from a Gaussian distributions with different standard deviations (*σ*). ***b***, *left*, variation (mean +/-std) of *k*−simplices vs *σ*. Inset shows relations at higher magnification. ***right***, number of cells (mean +/-std) in the target layer that receives suprathreshold (*n* ≥ *n*_*θ*_ = 5). Insets show representative synaptic fields. ***c***, same as above but with increasing number of active cells with a fixed *σ*

The relation between the number of functional simplices and the number of cells receiving suprathreshold inputs is not straightforward as there can be far more simplices in the source layer than neurons in the target layer (Fig. 4B). If the distance between active source neurons are within the dimension of a synaptic field (*λ* × *λ*), the number of possible combinations that a k-simplex that can be formed from *n* neurons is given by 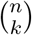. For the simulation in Fig. 4, the number of *k*-simplices with range from 100 (*k* = 1) to 3000 for (*k* = 6). In contrast, the number of cells receiving suprathreshold inputs ≥ *n*_*θ*_ = 5 increases to approximately 12 (B, right). Moreover, the number of active cells saturate while the higher dimension simplices continue to increase in an exponential manner (left).

### Categorization of spatial firing patterns for a single stimulus

The spatial distribution of neuronal firing will vary with stimulus conditions. To categorize the spatial patterns evoked under different conditions, we calculate the Betti numbers of the Čech complexes. The *n*-Betti numbers refers to the number of *n*-dimensional ‘holes’ in the topological space, thereby providing information about the local and global geometric relations between neurons.

The Betti numbers *β*_0_ and *β*_1_ represent, respectively, the number of 0 and 1 dimensional holes in the network of active neurons. Within the context of this study, *β*_0_ is the number of clusters whose neurons are functionally connected locally within but not across clusters. *β*_1_ is the number of ‘gaps’ within clusters where simplices are not contiguously connected, possibly indicating pockets of decreased activity in the target layer. These abstract concepts can visualized with the schematic in Figure 5, which shows the evolution of *β*_0_ and *β*_1_ as the number of active neurons is increased. With *n*_*θ*_ = 5 (*i*), there are 2 functional clusters of 4-simplices (tetrahedrons) that are separated so that *β*_0_ = 2. Adding a single point (*ii*, red), converts the triangles to tetrahedrons, effectively bridging the two clusters into one; thus, *β*_0_ = 1 and *β*_1_ = 0. Adding another point (*iii*), completes the ring of tetrahedrons, resulting in a hole in the middle so that *β*_1_ = 1. Finally, adding a point at the center (*iv*), fills the hole and again *β*_1_ = 0.

**Figure 5:**
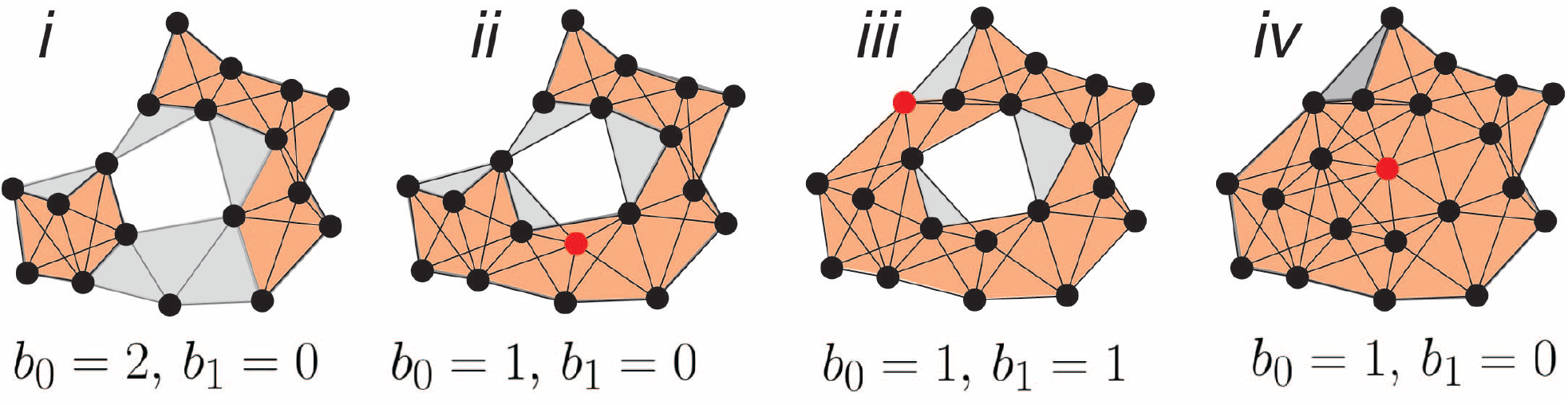
Evolution of Betti numbers. ***i-iv***, Vertices (red) are added sequentially to the Čech complex to illustrate changes in *β*_0_ and *β*_1_. Gray areas and lines represent non-functional simplices.

The higher Betti numbers for multidimensional holes do not have an obvious functional interpretation for 2 dimensional spaces. The cavities, given by *β*_2_, associated with hollow tetrahedrons are not counted because they do not exist in 2 dimensions when projected to a Euclidean plane (i.e. ‘flattened’). Betti number *β*_2_, however, would be relevant for a 3 dimensional network.

The evolution of *β*_0_ and *β*_1_ were examined by changing the spatial spread (*σ*) of a fixed number of source neurons (Fig. 6A, see above). At large spread *σ >* 6, the average *β*_0_ is greater than 1 (Fig. 6A*i*), indicating that there is variability in formation of 4-simplices from trial to trial. The average *β*_1_ is also around 2 (*ii*), indicating the presence of 1 dimensional holes within each cluster. At these large *σ* values, there are no suprathreshold inputs to target cells (gray curves). As *σ* decreases, *β*_0_ decreases to 1 while *β*_1_ reaches a peak and then becomes zero. The increase in density of neurons with decreasing *σ* induces high dimensional k-simplices, which in turn allow the 1-dimensional holes to form. However, with a further increase in density, the holes are filled as in Fig. 5. At this stage, the number of active target cells has increased substantially. (Fig. 6A*iii,iv*) show the evolution of *β*_2_ and *β*_3_. The evolution of the Betti numbers with increasing neuron density or number is consistent with that observed with increasing the number of edges in cliques in non-random networks [17]. Similar observations were made when the number of cells was increased at a fixed *σ* (B).

**Figure 6:**
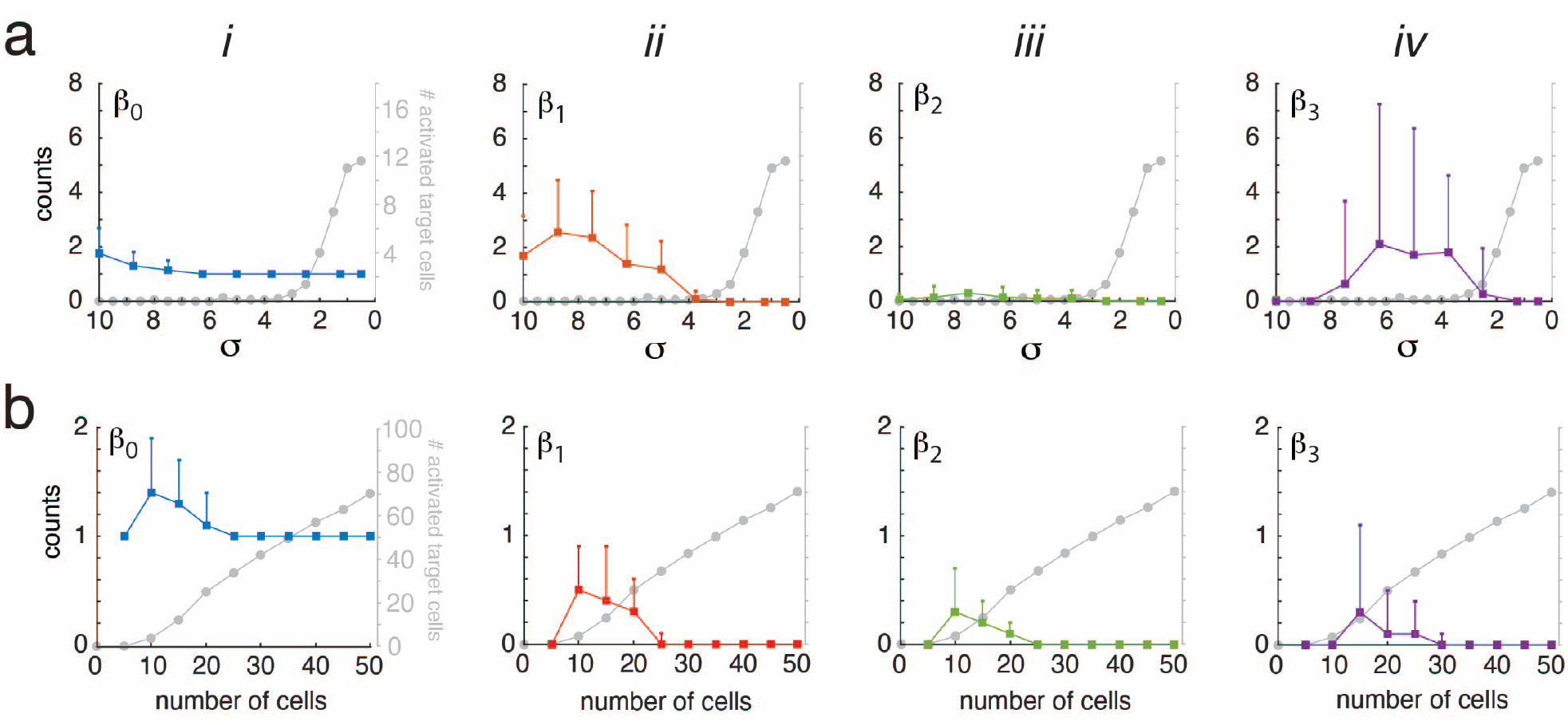
Betti numbers with changing stimuli. ***a, i-iv*** average (+/-std) Betti numbers *β*_0_ to *β*_3_ as a function of decreasing *σ* (constant number of cells). Superimposed gray traces show change in the number of target neurons receiving suprathreshold inputs. ***b***, Same as for ***a*** except that the number of source cells was increased with *σ* constant.

Comparing the evolution of the Betti numbers with the number of target neurons receiving suprathreshold inputs (gray curves) show that *β*_0_ and *β*_1_ were most variable when the the stimuli were weak (high *σ* (A) and few activated source cells (B)). When the stimuli became robust, *β*_0_ and *β*_1_ stabilized to 1 and 0, respectively.

### Distinguishing single from multiple activation patterns

Under natural conditions, multiple stimuli often occur simultaneously, possibly resulting in multiple clusters of activated neurons. Ambiguities in coding arise when the features of the stimulus differ by only a small amount, which in neural space may result in intermingling clusters (Fig. 1C). In the target layer, it becomes unclear whether the synaptic fields were due 1 or 2 clusters of active neurons in the source layer. In the following, we use Betti numbers to determine the number of active clusters during a complex stimulus.

Figure 7 shows 2 clouds of active neurons that are initially far apart so do not interact. When separated (left column) each cluster contains both functional connections (red) that generate suprathreshold events in the target layer (top) and non-functional connections (black in bottom, light blue in top). As the two clusters are brought closer together, additional connections (black) begin to form between clusters, though they remain non-functional (middle column). The functional simplices (red) remain unchanged and the overlap in the synaptic fields between the clusters appear but remain subthreshold. With a further decrease in distance, the two clusters merge into a single cluster with many more functional connections (right column). The overlap in synaptic fields increase substantially and more active areas emerge.

**Figure 7:**
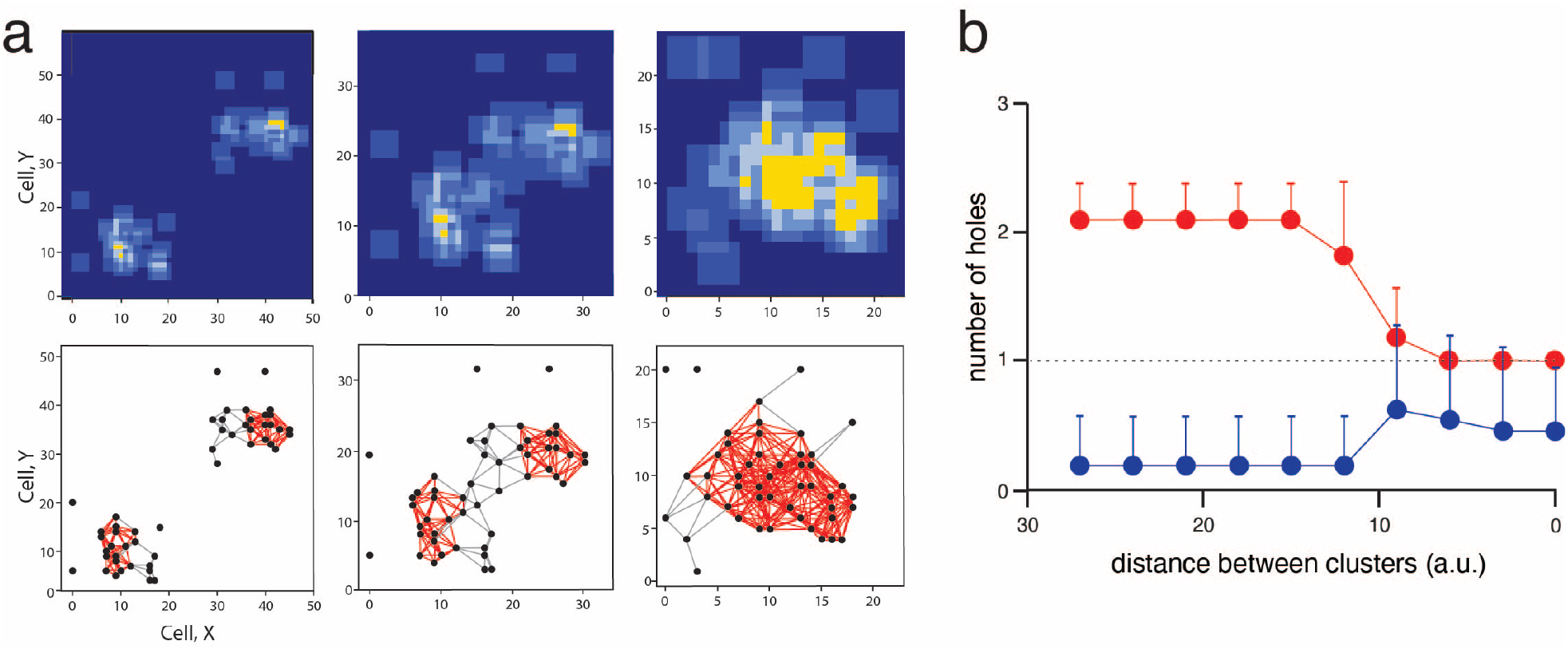
Discrimination of multiple clusters *columns 1-3*, two clusters of active source neurons were brought progressively close to each other. ***bottom***, Čech complexes consisting of 25 neurons each. ***top***, synaptic fields. Yellow show regions were neurons received suprathreshold input (*n*_*θ*_ = 5)

**Figure 8:**
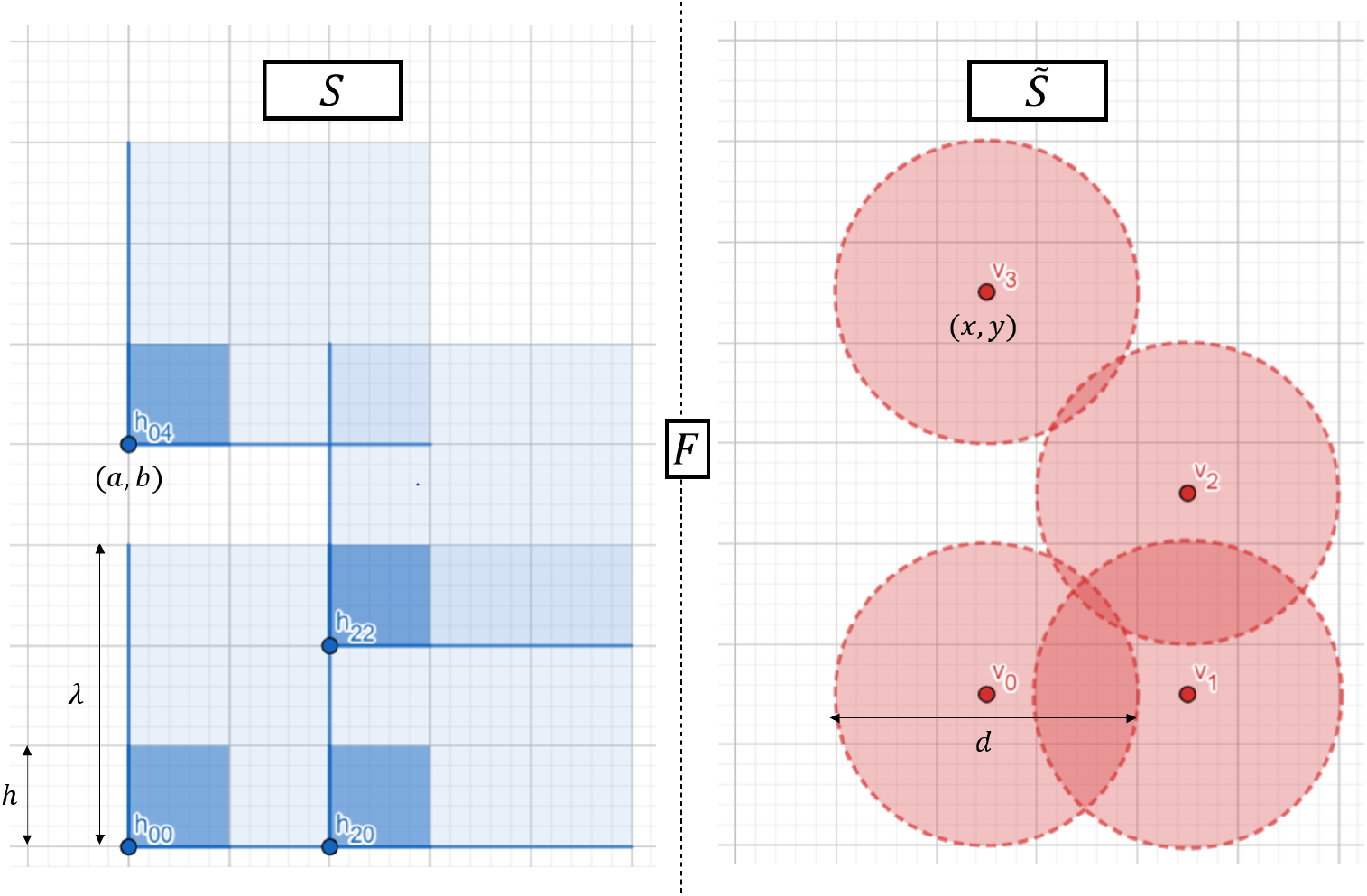
Graphical representation of the equivalence between spaces *S* and 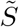

The changes in the connections are reflected in the Betti numbers. Fig. 7b shows the evolution of *β*_0_ and *β*_1_ (averaged over 50 trials) with decreasing distance between clusters. The *β*_0_ changes from 2 when the clusters are separated to 1, when the clusters are fully merged. Note that *β*_0_ = 2 even though the neurons in each cluster have begun to connect (distance = 15; A, middle column). The standard deviation is highest at the transition phase (distance between 10 and 15 units) because some of the neurons toward the edge of each cluster occasionally generate suprathreshold events. The *β*_1_ is low initially and then increases to a higher value when the clouds merge. In the transition phase, *β*_1_ increases to a peak before reaching a lower level. The rise is mediated by an increase in the total number of active neurons at the edge of each cluster and the fall is because the holes begin to be filled in. For the parameters of the simulation (25 neurons per cluster), higher Betti numbers are zero at all distances (not shown).

## Discussion

We developed a mathematical framework for identifying spatial patterns of activity in one layer that are necessary for propagation of signals across neural networks. We constructed Čech complexes in the source layer that are constrained by the extent of synaptic fields in the target layer. By specifying the minimum number of inputs needed for a target neuron to fire, the ‘functional’ simplices and the associated source neurons that form its vertices can be determined. This proved useful for distinguishing neurons within an active population that actually contribute to signal transmission and for detecting multiple clusters that may result from complex stimuli.

### Comparison to previous studies

Topological analysis [18] is commonly used to characterize the architecture of data sets. Typically, the Čech or Vietoris-Rips complexes are calculated by increasing the diameter of ‘balls’ and then constructing simplices and documenting the persistence of Betti numbers [19, 20], with the longer lasting Betti numbers being more indicative of the true topology. In this study, the size of the synaptic fields were kept constant and instead, the ‘cloud’ of active neurons was varied to reflect changes in e.g. the intensity of the stimulus [16]. The resultant Čech complexes changed accordingly, as did the Betti numbers. Stable Betti numbers *β*_0_ and *β*_1_ indicated when the source cells generated consistently threshold events in the target layer (Fig. 6) and when population activity transitioned between single and multiple clusters (Fig. 7).

Topology has also been used to elucidate coding schemes in neural networks. In hippocampus, the receptive fields of place cells were inferred based on whether the cells were active coherently within a short time period [21]. With the neurons as vertices, edges are drawn between pairs of co-firing neurons that presumably resulted from intersecting, albeit unspecified, receptive fields in stimulus space. Our approach is related except that the Čech complexes were computed by first specifying the extent of the synaptic fields in the target layer. In essence, our analysis applies to the next stage of processing. The spatial distribution of active source neurons presumably reflect the intersecting receptive fields originating in the sensory organs; because of topography, the activated cells will naturally be confined to an enclosed area. Our analysis reveals the subset of these stimulus-activated cells that are involved with propagation of information to the next stage of processing.

### Relation to temporal correlation

For this study, only the locations of active neurons in the source layer were important for determining whether their combined activity were sufficient to cause firing in the next layer. However, theory [5] and experiments [11, 12] indicate that synchrony is also important: jitter in the timing of action potentials can degrade propagation across multilayer networks.

The condition that the active neurons have to be both close to each other in space and synchronized is probably satisfied in topographically-organized networks. Neurons that are close to each other are likely to receive common external inputs and hence fire synchronously [22, 23, 24]. Moreover, topological analyses suggest that correlations between neurons is determined to a large extent by the architecture of the networks [17, 25]. Electrophysiological studies reveal that the connection probability between neurons are are Gaussianrather than uniformlydistributed [26]; hence, neurons that are close to each other in the source layer are more likely to be synaptically-connected and exhibit correlated activity. In principle, the formation of functional simplices could be modified to factor in both the extent of the synaptic field in the target layer and the correlated activity of neurons in the source layer.

### Functional implications

Understanding the conditions for when population activity in a network is propagated to another is of some significance. Given the small size of excitatory synaptic potentials in cortex (∼ 0.1-1 mV), several are needed to evoke action potentials in target neurons [27, 28, 29]. The lack of high dimension simplices in the Čech complex would signify that the activities are confined to a single layer. Hence, noise-related or spontaneous activity, even if synchronous, would not lead to firing in the next layer provided that the active neurons are spaced widely from each other. Alternatively, computations may be purposely confined to the layer by avoiding situations that lead to high dimension simplices.

The Betti numbers can be used to as a basis for initial categorization of spatial activity patterns. In the auditory system, for example, the location and size of the activated area may be used to encode the frequency and intensity of tones [16]. Psychophysical experiments show that the perceived frequency of a tone changes little with intensity [30]. The 0-Betti number (*β*_0_) is invariant to the size of the activated cluster and hence intensity. On the other hand, two tones are predicted activate separate clusters provided the frequency difference is sufficiently large; if not, the two areas will merge as one and the two tones will be perceived as a single tone [31, 32]. *β*_0_ would then change from 2 to 1, with the transition phase marked by large variability in both *β*_0_ and *β*_1_ as neurons at the edge of each cluster start to intermix.

The functional significance of *β*_1_ is less clear although there may be situations where there are gaps consisting of inactive neurons within a cluster of active neurons. For the scheme described here, the gaps arise primarily when e.g. the stimulus is weak and the active neurons in the source layer are sparse. However, there may be realistic scenarios where ‘holes’ that appear in a cluster have functional significance. In the auditory system, it is theoretically possible that the spatial distribution of neurons the source layer and the synaptic fields in the target layer are annuli: the axonal spread of some inhibitory neurons can be narrower than that of excitatory cells [26], the responses to two pure tones can produce response areas that have inhibition surrounded by excitation [33], and complex stimuli with a ‘spectral’ notch can produce a gap in the firing of neurons along the tonotopic axis [34].

## Limitations

The main advantage of the analysis is that general concepts can be proven using mathematical formalism. However, a major disadvantage is that several simplifications and assumptions were necessary. Consequently, there are several caveats when applying the analysis to biological systems.

The first is that synaptic fields are not perfect disks since axon collaterals diverge into several termination points (Fig. 1a, [3, 13, 14, 15]); the intersections and Čech complexes cannot be strictly computed. Under realistic conditions, the dimensions of simplices that lead to firing is likely to be underestimated, although it might be possible to introduce a correction term involving connection probability between neurons across layers.

The second is that there are other factors that determine neuronal firing that are not included in the analyses. Neurons that fire at high rates would generate temporally-summated synaptic potentials in target neurons thereby increasing the probability of firing and effectively reducing *n*_*θ*_. Spontaneous and/or background inputs to target cells would bring their resting potential closer to threshold. In both cases, the dimensions of simplices would be overestimated. To increase accuracy, the analysis could be restricted to brief stimuli.

The third is that the inhibitory neurons are not included in the analysis. Local inhibitory interneurons are common in biological networks [35, 26]. They are often co-activated by external input and will reduce the excitability of cells and cause an underestimate in the dimension of the threshold simplices. However, because there is often a delay [36, 37, 38] between the excitatory and inhibitory inputs, the analysis could be relatively unaffected if applied only for brief stimuli.

Finally, as mentioned in the Results, there is not a simple relationship between the number and dimension of simplices in the source layer and the number of target neurons receiving threshold inputs (Fig. 4). Similarly, the Čech complex cannot be used to estimate the size of the overlapping fields in the target layer (5). The strongest conclusion that can be drawn is that an *k*-simplex in the source layer means that at least one neuron in the source layer is receiving *k* + 1 convergent inputs. Conversely, a cell firing in the target layer means that there is at least one *n*_*θ*_ − 1 simplex in the source layer.

Despite these limitations, the theory is likely to be of some value for characterizing and categorizing data obtained with imaging techniques where the location of neurons are known (assuming that the extent of the synaptic fields can be estimated). Because precise timing is not crucial, the analysis will be particularly useful for calcium imaging where the signals are relatively slow [39, 40].

## Materials and Methods

### Generating 2D Gaussian-distributed activity patterns

A stimulus is assumed to increase the firing probability *G*(*x, y*) in neurons of the source layer *S*^*β*^. A discrete statistical distribution is needed because the location of cells (*x*_*i*_, *y*_*i*_) in neural space is determined by *x*_*i*_ ∈ χ = {*i* · *h* | 0 ≤ *i < N*_*x*_} and *y*_*i*_ ∈ 𝒴 = {*i* · *h* | 0 ≤ *i < N*_*y*_}. To control independently the number of cells and the standard deviation, we perform the following procedure.

We first create two arrays *X* = (*x*_0_, *x*_1_, …, *x*_*m*_) with *x*_*i*_ = *x*_min_ + *i* · *h*, and 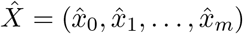, with 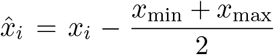 centered around zero. We then define Φ_*σ*_ (*z*_*i*_) = *ϕ*_*σ*_ (*Z* ≤ *z*_*i*_) as the continuous, one-dimensional gaussian cumulative distribution function for the random variable *Z* with standard deviation *σ* centered around *z* = 0. Every value *x*_*i*_ is then associated with the total probability accumulated in 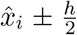 given by 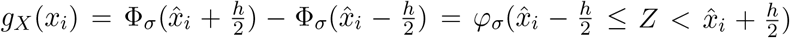. With this method, we are able to limit the extent of the distribution, produce a discretized Gaussian distribution with a specific standard deviation *σ*, and obtain a discretized probability vector *P*_*X*_. We obtain the probability vector *P*_*Y*_ for *Y* in a similar manner and combine the two probabilities by using the outer product of *P*_*X*_ and *P*_*Y*_. The distribution is normalized by dividing every value by the sum 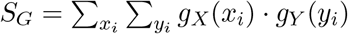. The resulting 2D probability is 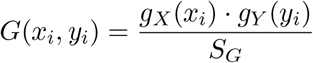

The above procedure yields a set *P* ⊂ χ × 𝒴 of possible points from the grid, each of which have an assigned probability. The final step is to specify the number of cells *N* for a given distribution. This is given by:

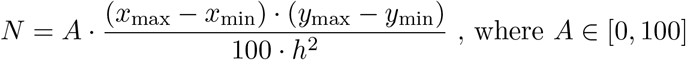

*N* non-repeated elements of the list *P* with probability *G*(*P*) are chosen randomly to obtain the set of points (i.e. active neurons) in the discrete space.

## APPENDIX

### Neural Space

#### Discretization into neurons

To facilitate analyses, the two dimensional neural space *S*^χ𝒴^ is defined as a product of two half-open intervals *S*^χ^, *S*^𝒴^:

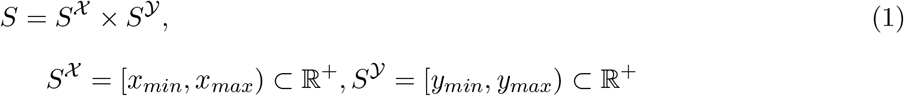

*S*^χ^ and *S*^𝒴^ are each partitioned into half-open sub-intervals [16]:

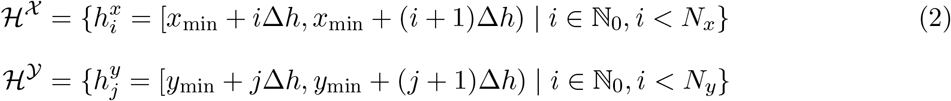

The sets containing the starting points of sub-intervals in both axes are:

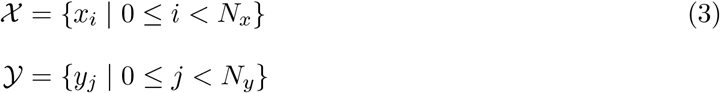

Visually, the neural space is a ‘half-open’ box with the bottom and left edges closed and the top and right edges open.

Similarly, individual neurons (Fig. 2A of Main text) are represented mathematically as ‘half-open boxes’ where the top/bottom and left/right sides are half open intervals *h*^*x*^ = [*x, x* + Δ*h*) and *h*^*y*^ = [*y, y* + Δ*h*), respectively. Because individual neurons are uniquely identified by the point at the lower, left point at the corner (henceforth termed ‘corner point’), they will be designated as *h*_*xy*_ = *h*^*x*^ × *h*^*y*^. The set of cells is then

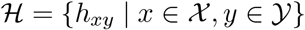

A topology *τ*_*S*_ for neural space *S* can be defined with ℋ as a basis.

##### Definition 1.

A topological space is a set χ together with a collection of open subsets *τ* that satisfies the conditions:

1. ∅, χ ∈ *τ*
2. The intersection of a finite number of sets in *τ* is also in *τ*.
3. The union of any number of sets in *τ* is also in *τ*.

Given the way the elements of the set ℋ are defined, we can check that ℋ is a basis of a topology because it satisfies the three conditions. Hence, the topology *τ*_*S*_ = {*h* ⊂ *S* | ∀*p* ∈ *h*, ∃*h*_*xy*_ ∈ ℋ s.t. *x* ∈ *h*_*xy*_ ⊆ *h*} is the topology generated by the basis ℋ and is a topology of the neural space *S*.

#### Synaptic field

The synaptic field in *S*^*β*^ (designated as *h*_*xyλ*_; *xy* ∈ χ ^*λ*^ ×𝒴^*λ*^) generated by a neuron in *S*^*α*^ is a half-open square that can encompass several neurons (eq. 1 of Main text). A mapping from neurons in *S*^*α*^ to synaptic fields in *S*^*β*^ is defined as (eq. 2 of Main text):

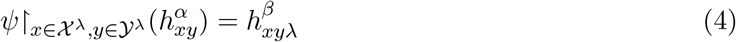

##### Lemma 1.

The mapping *ψ* from individual neurons the source space (*S*^*α*^) to synaptic fields in the target space (*S*^*β*^) is an homomorphism when the coordinates of the active cells are restricted 𝒜 ⊂ χ ^*λ*^ ×𝒴^*λ*^.

*Proof*. Because cells in the source layer are either the same or do not intersect at all, *S*^*α*^ can be equipped with the union operation of individual elements in the source layer:

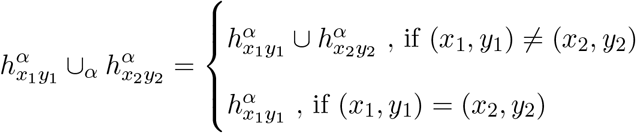

Then, let *S*^*β*^ be equipped with the union of two-dimensional intervals

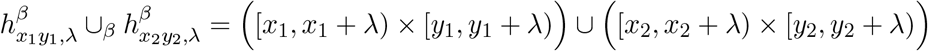

Hence, for every pair of different elements of 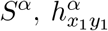 and 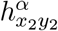 restricted to 𝒜 ⊂ χ ^*λ*^ × 𝒴^*λ*,^ the mapping *ψ* ↾ _A_ trivially preserves the union operation:

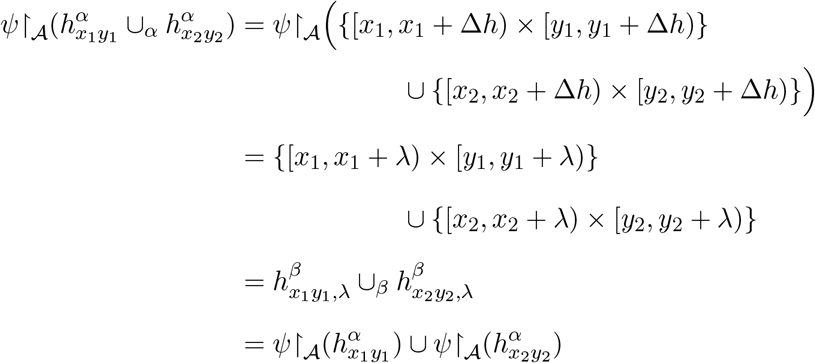

And therefore it is an homomorphism.

### Calculation of Čech Complex

#### Equivalence between rectangular synaptic fields and open balls

For the discretized neural space *S*, individual neurons *h*_*xy*_ and synaptic fields *h*_*xyλ*_ are half-open boxes in a plane, with synaptic fields identified by (*x, y*) and encompassing a contiguous set of *n*_*λ*_ neurons. In the following, we show that the rectangular synaptic fields can be made equivalent to a space 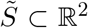 where the synaptic fields are balls *B*_*r*_(*v*_*i*_) centered around each cell in the source layer.

Since the simplicial complex is uniquely determined by the distance between vertices, we must prove that the generated simplicial complexes in *S* and in 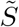 are the same. This is equivalent to the condition that synaptic fields 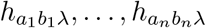 intersect in *S* if, and only if, balls *B*_*r*_(*v*_1_), …, *B*_*r*_(*v*_*n*_) intersect in 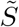. The relations that must be specified are between the points representing cells and the extent of the fields.

##### Theorem 1.

Let *S* ⊂ ℝ ^2^ be a neural space that is partitioned into individual square neurons each of width *h* and have square synaptic fields with corner points (*a, b*) = (*n*_*a*_ · *h, n*_*b*_ · *h*) ∈ *S* ⊂ ℝ ^2^ of length *λ* = *n*_*λ*_ · *h* with *n*_*a*_, *n*_*b*_, *n*_*λ*_ ∈ ℤ, *n*_*λ*_ ≠ 0 and *h* ∈ ℝ ^+^. Now define 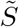 as a space consisting of balls with centers located at 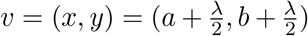 with fixed radii 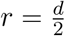 such that 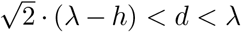. Then, ∀*I* = {*i*_1_, …, *i*_*n*_}, synaptic fields 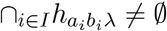 in 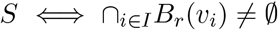 in 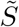

*Proof*. We will prove the two implications by stating which conditions need to be satisfied for the synaptic fields to intersect, taking into account that *S* is discretized in steps of size *h*.

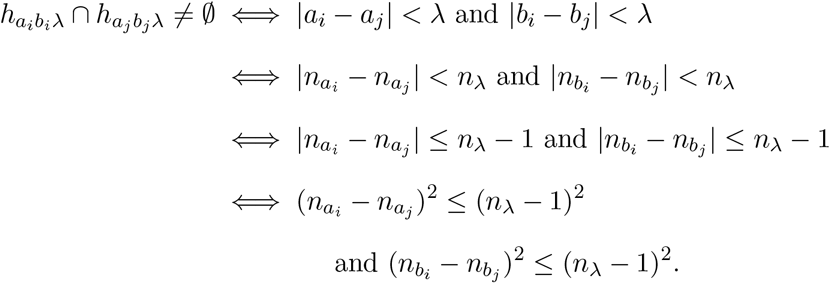

Therefore, in the limit case 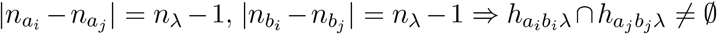. However, in particular, either 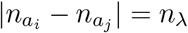 or 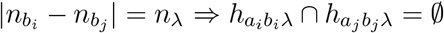.

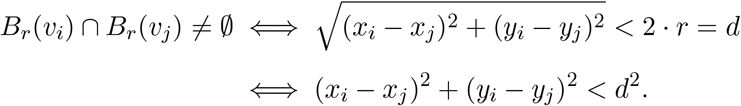

We will prove the two implications of the statement individually.

- 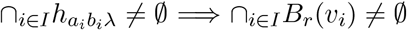

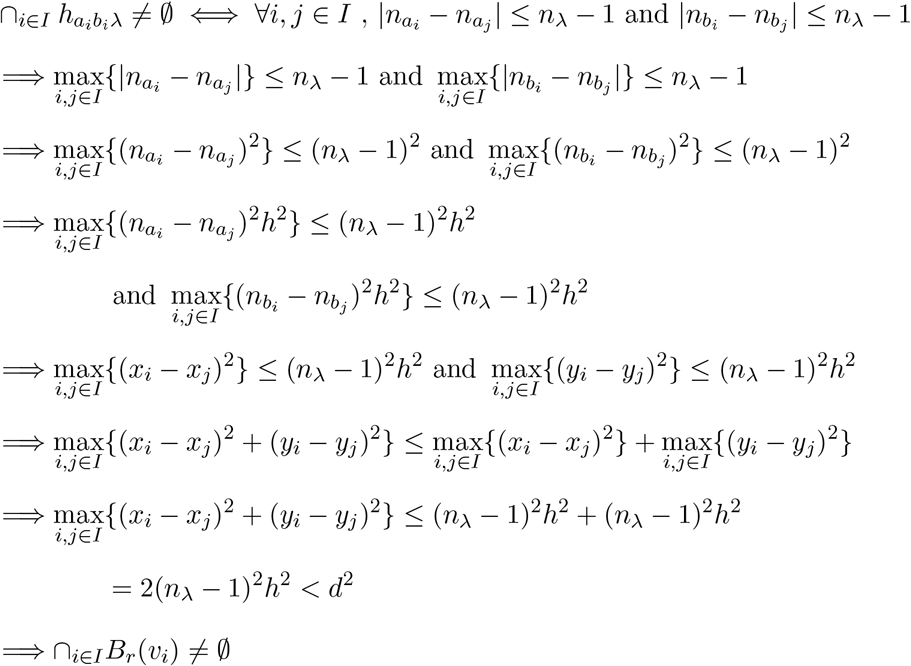
- 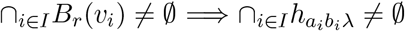

We can prove this using the contrary statement: instead of proving A ⇒ B, we prove that not(B) ⇒ not(A). By logic reasoning, it is equivalent to proving that 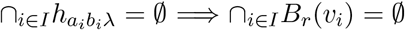.

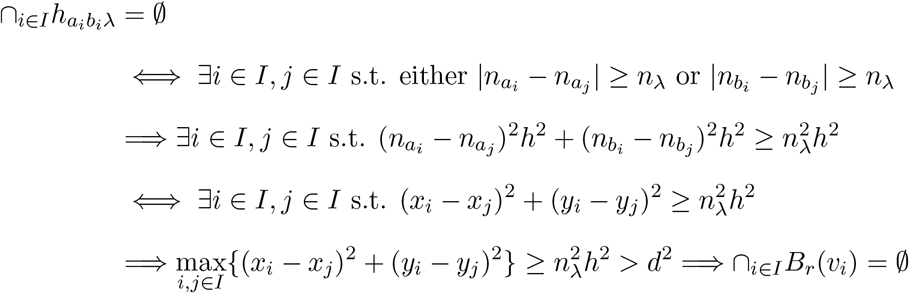

Therefore, 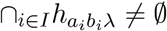 in 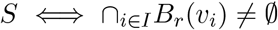 in 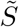.

#### Keeping track of coordinates of vertices in neural space

Although the abstract simplicies facilitate visualization of the network geometry, it will later be necessary to keep track of the coordinates of corner points that the vertices in *Č*(𝒜) represent. We thus define a graph 𝒢, with vertices *v* ∈ 𝒱 and edges *e* ∈ ε with a bijective mapping:

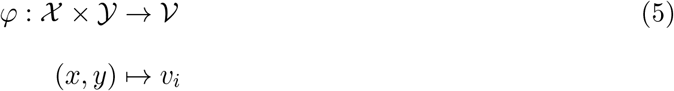

Consequently, the set of corner points 𝒜 ⊂ χ ×𝒴 can be related directly to the vertices in 𝒱: 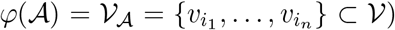, with the set of edges 𝒱corresponding to ‘connections’ between corner points. The simplices of Č in abstract space can then be converted to simplices in Euclidean neural space.

#### Functional simplices

Because neurons are tuned to a particular feature of the stimulus and are topographically-organized, the set of activated presynaptic cells are clustered within a confined area. Consequently, set of synaptic fields in the target network will also be confined and will overlap, depending on the size of *λ*. To a first approximation, the amount of input a target cell firing is determined by the degree of overlap in the synaptic fields. Neurons will fire if the number of inputs exceed a threshold value *n*_*θ*_. A set of synaptic fields intersecting with each other can be defined as 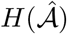 :

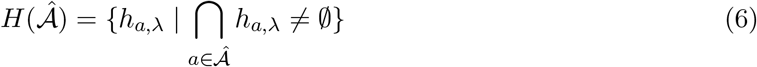

A postsynaptic cell in the target network will fire if 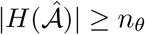, where *n*_*θ*_ is the number of inputs needed to cross threshold. We show below that this requirement is met when the presynaptic neurons in the source network forms a simplex with ≥ *n*_*θ*_ vertices and, therefore, the set of synaptic fields that are part of the functional network is *H*^*θ*^.

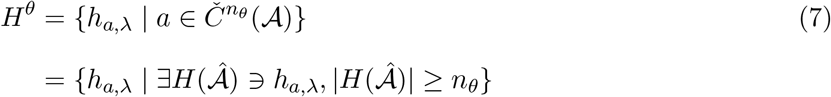

We show below that this requirement is met when the presynaptic neurons in the source network forms a simplex with *n*_*θ*_ or more vertices. First, we show that:

##### Lemma 2.

The set *H*^*syn*^ consisting of synaptic intervals with dimensions *λ*_*x*_ × *λ*_*y*_ forms an open cover for *S* and specifically for the set of corner points of synaptic boxes in *H*^*θ*^ ({(*x, y*) ∈ *C*_*xy*_ | *h*_*xyλ*_ ∈ *H*^*θ*^}).

*Proof*. By construction, neural space *S* is partitioned into cells consisting of half-open boxes. Hence, the set of cells is a cover for *S*:

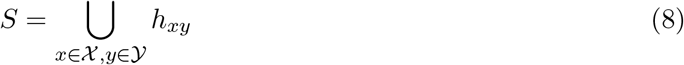

Similarly, a synaptic area is a union of contiguous sets of cells:

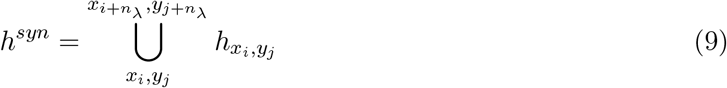

Let the sets containing the coordinates of the synaptic intervals be χ ^*λ*^ = {0, 1, …, *N*_*x*_−*λ*}, 𝒴^*λ*^ = {0, 1, …, *N*_*y*_ − *λ*}. Then:

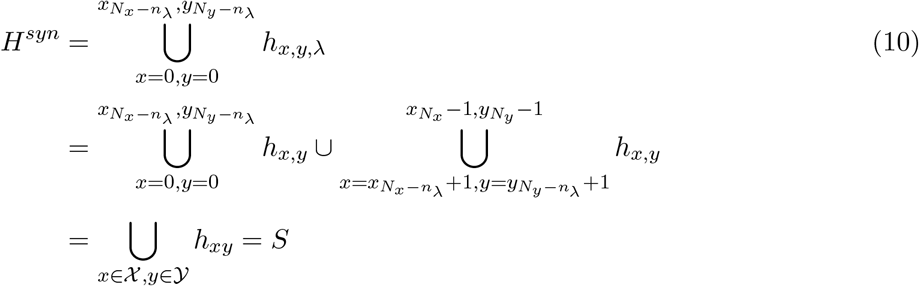

since χ = χ ^*λ*^ ∪ {*N*_*x*_ − *n*_*λ*_ + 1, …, *N*_*x*_ − 1}, 𝒴 = 𝒴^*λ*^ ∪ {*N*_*y*_ − *n*_*λ*_ + 1, …, *N*_*y*_ − 1} and so is a cover. Since *H*^*syn*^ ⊂ *τ*_*S*_, it is also an open cover. Finally, since each synaptic box contains the corner point, the synaptic boxes are automatically covers for corner points of synaptic boxes in *H*^*θ*^

##### Theorem 2.

*Proof*. Letting *P* = {*p*_*k*_ = (*x*_*k*_, *y*_*k*_) ∈ χ^*λ*^ ×𝒴^*λ*^}, *B*_*λ*_(*p*_*k*_) be a half-open box with edge length of *λ* with closed corner (lower left corner) *p*_*k*_, and *σ* be a simplex, a Čech complex in *S*^*β*^ is then defined as:

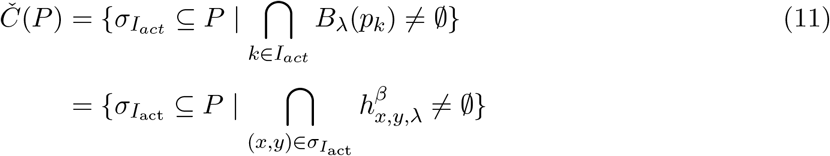

If a 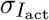 of the set is a simplex with dimension ≥ (*n*_*θ*_ − 1), then *n*_*θ*_ vertices, or neurons 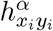, in the source layer are connected and the intersection of their respective synaptic fields 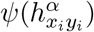 in *S*^*β*^ is not null. Therefore, neurons 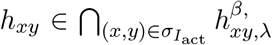 received sufficient inputs to fire action potentials. Therefore, if one of the elements of the set is a simplex of dimension at least (*n*_*θ*_ − 1), there is a region in *S*^*β*^ that receives suprathreshold input and any neuron in that region will fire an action potential.

In *S*^*α*^, we construct a Čech complex with cells of length *ϵ*. Note that in *S*^*β*^ the separation between corner points of each element of the set have Chebyshev distances less or equal than *λ*. The pre-image (*ψ*^−1^(*H*^*θ*^)) is a set of points in *S*^*α*^, which would be the positions of the cells that originate those fields and, therefore, the cells in *S*^*α*^ that form the Čech complex.

Because of the bijective mapping defined above, the points *p* separated by distances less than a certain *λ* in the Čech complex in *S*^*β*^ correspond to points 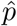 in a Čech complex in *S*^*α*^ using 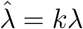, with *k* ∈ ℝ^+^ a scaling factor.

#### Algorithm for computing Čech complexes

The task is to calculate the abstract simplicial complex Č given the parameters *λ, n*_*θ*_ and a set of cells 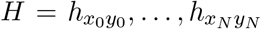, each of which has an assigned position (*x, y*). Č is computationally described by *C* = {*σ*_0_, …, *σ*_*m*_} where every element of this list is a short array *σ*_*i*_ = [*i*_0_, …, *i*_*k*_] that contains the indices of a set of (*k* + 1) cells from *H* that form a *k*−simplex in Č.

In this discretized space with square synaptic fields, determining if a set of *k* + 1 cells that forms a *k*−simplex involves calculating the Chebyshev distances between points from the set in the *x* and *y* axis and checking if all of them are smaller than the fixed extent of a synaptic field *λ*.

##### Determining all the simplices of the complex

Computationally, calculating the distances between all pairs of *k* points from a set *H*_*I*_ is a *O*(*k*^2^) problem without any possibility for a better solution. However, the complexity may be lowered to *O*(*k* · log (*k*)) by directly discarding any distance that is greater or equal to *λ* since is not a valid connection between cells. The points are sorted lexicographically, first by *x* and then by *y* and checking the difference between the first and last value of the sorted list in every respective case. We then check whether every nonempty subset of *k* cells in *H*, for 0 *< k* ≤ *N* form a simplex. The total number of nonempty subsets in a finite set of *N* elements, including the whole set, is 2^*N*^ − 1. However, since each cell is a 0-simplex, the total is reduced to 2^*N*^ − *N* − 1.

Calculating every possible simplex in the Čech complex increases the complexity of the task substantially. However, because we are interested in the simplices and holes of dimension up to not much more than *n*_*θ*_, it is possible to limit the dimension of the simplices up to a threshold *M* ; *n*_*θ*_ *< M* ≤ *n*_*θ*_ + 3. Simulations indicate that this is sufficient to reveal the main properties of the resulting networks. Given that the total number of different nonempty subsets of dimension *k* ≥ 2 from a finite set of *N* elements is 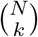, the number of times it would be then necessary to check if the given subsets forms a simplex or not is 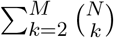.

The Čech complex can be constructed efficiently by considering only the maximal simplices, which are simplices that cannot be extended to a valid simplex (according to our maximum edge length) by adding an extra point. Further improvements can be made to the algorithm by starting with sets of higher dimension *k* = *M* and lowering the dimension until *k* = 1 calculating every maximal simplex at each step, so that the calculation does not have to be repeated for every subset of simplices with lower dimensions. For example, if at *k* = 4 we find that the set of cells {*a, b, c, d*} forms a 3−simplex, then we will also add the 2−simplices {*a, b, c*}, {*a, b, d*}, {*a, c, d*}, {*b, c, d*} the 1−simplices {*a, b*}, {*a, c*}, {*a, d*}, {*b, c*}, {*b, d*}, {*c, d*} and the 0−simplices {*a*}, {*b*}, {*c*}, {*d*} to the abstract simplicial complex, and discard the possibility of having to check all those combinations again in the lower-dimension steps.

Once we have the abstract simplicial complex, we can add every simplex to a `Simplex Tree` object from the `Python Gudhi` library, a task of order *O*(*m*), where *m* is the total number of simplices in Č. This object is very useful because it already has the internal homology methods to calculate the Betti numbers of the generated complex.

